# Adipokines contribute to central-obesity related reductions in myelin-sensitive MRI indices in the fornix

**DOI:** 10.1101/440990

**Authors:** Claudia Metzler-Baddeley, Jilu P. Mole, Erika Leonaviciute, Rebecca Sims, Emma J. Kidd, Benjamin Ertefai, Aurora Kelso-Mitchell, Florence Gidney, Fabrizio Fasano, John Evans, Derek K Jones, Roland J. Baddeley

**Affiliations:** Cardiff University Brain Research Imaging Centre (CUBRIC), School of Psychology, Cardiff University, Maindy Road, Cathays, Cardiff, CF24 4HQ, UK; Psychological Medicine and Clinical Neurosciences, School of Medicine, Cardiff University, Maindy Road, Cathays, Cardiff, CF24 4HQ, UK; School of Pharmacy and Pharmaceutical Sciences, Cardiff University, Redwood building, King Edward VII Avenue, Cardiff, CF10 3NB; Siemens Healthcare, Head Office, Sir William Siemens Square, Surrey, GU16 8QD, UK; School of Psychology, Faculty of Health Sciences, Australian Catholic University, Melbourne, Victoria 3065, Australia; Experimental Psychology, University of Bristol, 12a Priory Road, BS8 1TU, UK.

## Abstract

Midlife obesity is a risk factor of late onset Alzheimer’s disease (LOAD) but why this is the case remains unknown. As systemic inflammation is involved in both conditions, one possibility is that obesity-related neuroinflammation may contribute to the development of LOAD. Neuroinflammation is closely linked to white matter myelin loss, and this can be measured *in vivo* with quantitative magnetization transfer (qMT) imaging. Here, we investigated whether differences in obesity measures, i.e., in Waist Hip Ratio (WHR), abdominal visceral and subcutaneous fat volume fractions and Body Mass Index (BMI), were associated with reductions in qMT indices of apparent myelin in temporal white matter pathways involved in LOAD (i.e., the fornix, the parahippocampal cingulum and the uncinate fasciculus compared with whole brain and cortico-spinal white matter) in 166 cognitively healthy individuals (38-71 years of age). Obesity-related effects on myelin-sensitive markers were contrasted with differences in apparent axon density from dual-shell diffusion Neurite Orientation Dispersion and Density Imaging (NODDI). Differences in WHR and in visceral fat volume fractions were negatively correlated with differences in qMT estimates of apparent myelin and tissue metabolism in the fornix but not with any other microstructural components. These correlations were not explained by demographic (age, sex, education), health (hypertension, alcohol consumption, sedentary lifestyle) or genetic (*APOE* genotype, family history of dementia) risk factors of LOAD. Furthermore, differences in the ratio of plasma concentrations of leptin and adiponectin were also positively correlated with differences in C-Reactive Protein concentrations, and contributed significantly to the correlations between central obesity and myelin-sensitive metrics in the fornix. These results are consistent with the view that visceral fat-related chronic inflammation may damage white matter myelin in limbic regions, known to be vulnerable to LOAD pathology.

## Introduction

Obesity is globally on the rise (WHO, 2018), and has become an epidemic in many Western countries. In the UK, two-thirds of adults are overweight or obese, defined by a Body Mass Index (BMI) of > 25kg/m^2^ or 30kg/m^2^ respectively. Western-style diet and sedentary lifestyles contribute to obesity risk and obesity-related diseases including metabolic syndrome, type 2 diabetes, and cardiovascular disease (Cox et al., 2015). Several epidemiological studies have also identified a positive association between midlife obesity and the incidence of late onset Alzheimer’s disease (LOAD) (estimated risk ratio of ~1.4) (Beydoun et al., 2008; Pedditizi et al., 2016). While the effects of excessive adiposity are complex and involve multiple immune, metabolic, and endocrine factors, it is increasingly recognised that persistent, low-grade inflammation may play a key role in obesity and may also contribute to the development of obesity-related diseases such as LOAD (Angelova and Brown, 2015; Bartzokis, 2011; Brown, 2009; Conde and Streit, 2006; Heneka et al., 2015; Papuc and Rejdak, 2017; Sochocka et al., 2018).

Neuroinflammation can lead to white matter myelin damage (Di Penta et al., 2013; Pang et al., 2010; Serres et al., 2009b), that may occur independently of axonal injury (Bitsch et al., 2000). It is possible to quantify apparent white matter myelin changes *in vivo* with quantitative magnetization transfer (qMT) (Ou et al., 2009; Schmierer et al., 2007; Sled, 2017; Weiskopf et al., 2013; Whitaker et al., 2016), an MRI technique that is sensitive to changes in macromolecular density. The relative signal fractions from free water and semisolid macromolecular constituents of tissue are estimated in qMT by applying off-resonance radiofrequency pulses with constant amplitudes that are varied between frequency offsets. This selectively saturates the macromolecular magnetization with subsequent exchange processes resulting in magnetization transfer between saturated macromolecules and free water (Sled, 2017). In white matter, magnetization transfer is dominated by myelin (Ceckler et al., 1992; Koenig, 1991), and is also sensitive to microglia-mediated inflammation (Levesque et al., 2010; Schmierer et al., 2007; Serres et al., 2009a). The relative number of protons in the macromolecular pool, the macromolecular proton fraction (MPF), provides an index of apparent myelin content of white matter (Figure 1A). The rate of the magnetization transfer process *k*_*f*_ (Sled, 2017) has been shown to be sensitive to acute neuroinflammation in response to typhoid vaccination (Harrison et al., 2015), and has been proposed to reflect metabolic efficiency of mitochondrial function (Giulietti et al., 2012).

Here we applied MPF and *k*_*f*_ to investigate the hypothesis that central obesity is associated with reductions in apparent myelin/tissue metabolism of brain white matter in 166 cognitively healthy individuals between 38 and 71 years of age from the Cardiff Aging and Dementia Risk Study (CARDS).

QMT estimates of apparent white matter myelin were contrasted with MRI estimates of axon microstructure from multi-compartment diffusion based neurite orientation dispersion and density imaging (NODDI) (Zhang et al., 2012). NODDI provides separate indices of apparent axon density [intra-cellular signal fraction (ICSF)], of neurite orientation dispersion (OD), and of tissue free water contamination [isotropic signal fraction (ISOSF)] (Figure 1A). Thus, the combination of qMT and NODDI indices allowed us to study whether obesity-related differences in white matter microstructure were due to changes in apparent glia myelin/metabolism and/or apparent axon density/orientation.

Most imaging studies into obesity have investigated differences in BMI, an index that does not capture variation in body fat distributions (Adab et al., 2018). However, there is increasing recognition that it is not body fat *per se* but visceral rather than subcutaneous fat which leads to adverse health effects and increased risk of metabolic syndrome and mortality (Koster et al., 2015; Koster et al., 2010). Here, we therefore assessed obesity not only with BMI, but also with metrics of central obesity, i.e., the Waist Hip Ratio (WHR) and MRI indices of visceral fat volume fractions (viscVF) and subcutaneous fat volume fractions (subcVF) (Figure 1C). The examples in Figure 1C demonstrate the considerable individual variation in the distribution of abdominal subcutaneous and visceral fat.

As midlife obesity, notably visceral abdominal obesity, is a risk factor of LOAD, and LOAD pathology is known to spread from the hippocampal formation *via* limbic white matter pathways such as the fornix (Plowey and Ziskin, 2016) to the neocortex (Braak and Del Trecidi, 2015), we hypothesised that obesity-related changes would disproportionally affect limbic white and gray matter (fornix, parahippocampal cingulum, uncinate fasciculus, hippocampus) relative to whole brain and corticospinal-motor white matter (Kullmann et al., 2015; Metzler-Baddeley et al., 2013) (Figure 1B).

**Figure 1.**
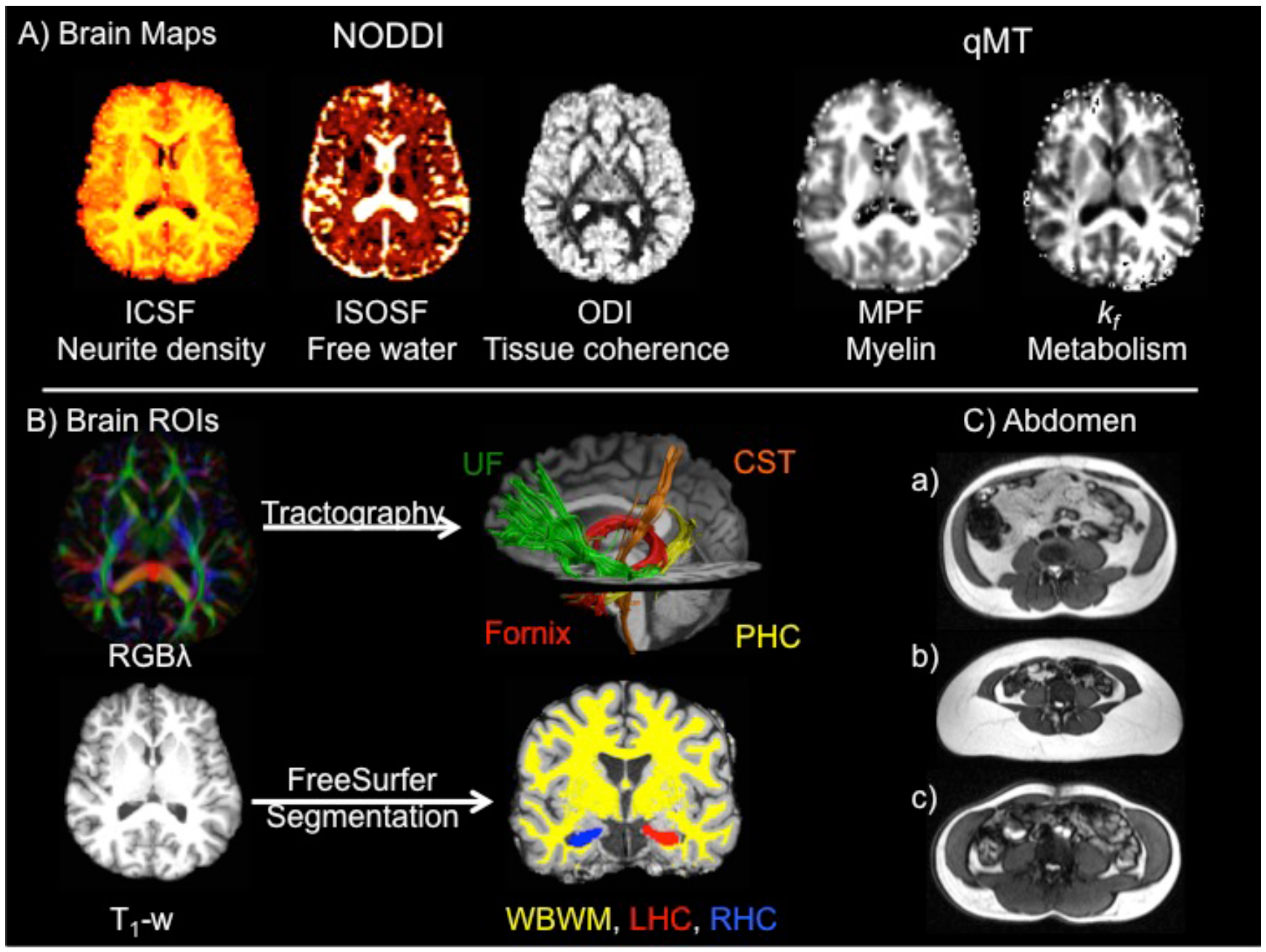
A) displays the MRI modalities and maps acquired from dual-shell high angular resolution imaging (HARDI) and from quantitative magnetization transfer (qMT) imaging. HARDI data were modelled with neurite orientation dispersion and density (NODDI) yielding maps of intracellular signal fraction (ICSF), isotropic signal fraction (ISOSF) and orientation density index (ODI). qMT based maps were the macromolecular proton fraction (MPF) and the forward exchange rate *k*_*f*_. B) Mean indices of the metrics were extracted from the left hippocampus (red), right hippocampus (blue), whole brain white matter (WBWM) mask (yellow), fornix (red), parahippocampal cinguli (PHC) (yellow), uncinate fasciculi (UF) (green) and corticospinal tract (CST) (orange). Hippocampi and WBWW were segmented from T_1_- weighted images with FreeSurfer version 5.3 and fornix, PHC, UF and CST were reconstructed with damped-Richardson Lucy spherical deconvolution (dRL) based deterministic tractography on colour coded principal direction maps (RGBλ). C) Examples of abdominal images from individuals with a) larger visceral than subcutaneous fat volume fraction, b) larger subcutaneous than visceral fat volume fraction and c) low visceral and subcutaneous fat volume fractions.

Mean values of all qMT and NODDI MRI indices were extracted from all regions of interest. Fornix, left and right parahippocampal cinguli, uncinate fasciculi, and corticospinal tracts were reconstructed with spherical deconvolution-based deterministic tractography in ExploreDTI (version 4.8.3) (Leemans et al., 2009) (Figure 1B). Whole brain white matter and bilateral hippocampal masks were segmented with FreeSurfer (version 5.3) (Fischl et al., 2002; Han and Fischl, 2007) (Figure 1B). The hippocampal regions included areas of the presubiculum, subiculum, cornu ammonis subfields 1-4, dentate gyrus, hippocampal tail and fissure (Iglesias et al., 2015; Iglesias et al., 2016) (Figure 1B).

**Table 1.** Summary of participants’ demographic, cognitive, anthropomorphic, physiological and genetic information

| Mean (SD) | Participants (n = 166) |
| --- | --- |
| Age (in years) | 55.8 (8.2) |
| Females | 56% |
| Years of education | 16.5 (3.3) |
| NART | 116.7 (6.7) |
| MMSE | 29.1 (1.0) |
| Positive Family History dementia | 35.1% |
| *APOE* Genotype | 39.9% $\epsilon 4+$, 60.1% $\epsilon 4-$ |
| Central obesity (Waist Hip Ratio) | 61.3% |
| Overweight/Obese (BMI) | 63.9% |
| Subcutaneous volume fraction | 0.39 (0.1) |
| Visceral volume fraction | 0.2 (0.06) |
| Systolic Hypertension | 28% |
| Smokers | 5.4% |
| Diabetes | 1.8% |
| Statins | 7.2% |
| Alcohol units per week | 7.4 (9.3) |
| Physical activities<sup>*</sup> | 10.3 (12.1) |
| (Median hours per week) |  |
| Adiponectin log<sub>10</sub> ng/ml | 4.04 (0.26) |
| Leptin log<sub>10</sub> pg/ml | 4.05 (0.47) |
| C-Reactive Protein log<sub>10</sub> ng/ml | 3.05 (0.47) |
| Interleukin-6 log<sub>10</sub> pg/ml | 0.7 (0.15) |

To disentangle effects of central adiposity from other health-related variables and to study the potential link with risk factors of LOAD, information about education, hypertension, alcohol consumption, physical activity, *APOE* genotype and family history of dementia were also collected (Dommermuth and Ewing, 2018; Ricci et al., 2017) (Table 1). Furthermore, we measured plasma concentrations of high sensitivity C-Reactive Protein (CRP) and interleukin-8 as markers of systemic inflammation (Swardfager et al. 2010) as well as of leptin and adiponectin, two adipokines that are involved in glucose control and the modulation of inflammatory responses (Arnoldussen et al., 2014; Doherty, 2011; Farr et al., 2006; Flak and Myers, 2016; Myers et al., 2008; Ryan et al., 2003). Leptin is known to up-regulate pro-inflammatory cytokines, while adiponectin has anti-inflammatory properties and down-regulates the release and the expression of pro-inflammatory cytokines (Lopez-Jaramillo et al., 2014).

Inter-individual differences in central obesity, specifically in visceral abdominal fat were expected to be associated with differences in MRI markers of apparent myelin/inflammation (MPF and *k*_*f*_) in limbic white matter pathways, notably the fornix and PHC. We also expected obesity to be accompanied by increases in ISOSF, a marker of CSF partial volume that may reflect unspecific tissue loss. However, we did not expect obesity to reduce the density of axons (ICSF) or alter their orientation dispersion in white matter. The application of the microstructural metrics to the hippocampus should be seen as exploratory, as microstructural indices are more difficult to interpret in gray relative to white matter due to its more complex structure. For this reason, and to test our assumption that qMT and NODDI indices would provide separable measurements of uncorrelated tissue properties, we also explored their dimensionality in white and gray matter regions separately, with principal component analyses.

## Methods

### Participants

Over a period of three years (2014-2017), n = 211 community-dwelling individuals between 35 and 75 years of age were recruited for CARDS from local Cardiff University databases and *via* internet and poster advertisements. The study was approved by the School of Psychology Research Ethics Committee at Cardiff University (EC.14.09.09.3843R2) and all participants gave written informed consent in accordance with the Declaration of Helsinki. Exclusion criteria were a history of neurological disease (e.g. Multiple Sclerosis, Parkinson’s disease, Huntington’s disease), psychiatric disease (e.g. schizophrenia, bipolar disorder, depression requiring hospitalization or a current PHQ-9 score of > 15 indicating severe depression), moderate to severe head injury with loss of consciousness, drug or alcohol dependency, high risk cardio-embolic source (mitral or severe aortic stenosis, severe heart failure, cardiac aneurysm), known significant large-vessel disease (i.e. more than 50% stenosis of carotid or vertebral artery, known peripheral vascular disease, coronary bypass or angioplasty) and MRI contraindications (e.g. pacemaker, cochlear implants, metal pins, stents, screws etc.). Demographic and health information including information about genetic and lifestyle risk factors of dementia was collected for all 211 volunteers. Here we report data from n = 166 who also underwent MRI scanning at the Cardiff University Brain Research Imaging Centre (CUBRIC). Table 1 provides a summary of the demographic, health and genetic information available for these 166 participants.

### Assessment of body composition/adiposity

Abdominal adiposity was assessed by measuring participants’ waist and hip circumferences to calculate WHR following the World Health Organisation’s recommended protocol (Organisation, 2008). Abdominal obesity was defined as a WHR ≥ 0.9 for males and ≥ 0.85 for females. BMI was calculated from participants’ height and weight. Normal weight was defined as a BMI of 18-24.9 kg/m^2^, overweight as BMI of 25-29.9kg/m^2^ and obese as BMI > 30kg/m^2^. Abdominal and subcutaneous fat volume fractions were obtained from MRI segmentation as described below. Systolic and diastolic blood pressure (BP) was measured with a digital blood pressure monitor (Model UA-631; A&D Medical, Tokyo, Japan) whilst participants were comfortably seated with their arm supported on a pillow. The average of three BP readings was taken and hypertension was defined as systolic BP ≥ 140 mm Hg. Other cardio-vascular risk factors of diabetes mellitus, high levels of blood cholesterol controlled with statin medication, history of smoking and weekly alcohol intake were self-reported by participants in a medical history questionnaire (Metzler-Baddeley et al., 2013). Information about participants’ physical activity over the preceding week was collected with the short version of the International Physical Activity Questionnaire (IPAQ) (Craig et al., 2003). The median number of hours of non-sedentary activities including walking, gardening, housework and moderate to vigorous activities were recorded. Participants’ intellectual function was assessed with the National Adult Reading Test (NART) (Nelson, 1991), cognitive impairment was screened for with the Mini Mental State Exam (MMSE) (Folstein et al., 1975) and depression with the Patient Health Questionnaire for Depression (PHQ-9) (Kroenke et al., 2001). All participants were cognitively healthy and scored at superior level of intelligence in the NART. Eight participants scored ≥ 10 in the PHQ-9 suggesting moderate levels of depression but no participant was severely depressed.

#### Blood plasma analysis

Venous fasting blood samples were drawn into 9ml heparin coated plasma tubes after 12 hours overnight fasting and were centrifuged for 10 minutes at 2,000xg within 60 minutes from blood collection. Plasma samples were transferred into 0.5 ml polypropylene microtubes and stored in a freezer at −80°C.

Circulating levels of high-sensitivity CRP in mg/dL were assayed using a human CRP Quantikine enzyme-linked immunosorbent assay (ELISA) kit (R & D Systems) and interleukin-8 levels in pg/mL were determined using a high sensitivity CXCL8/INTERLEUKIN-8 Quantikine ELISA kit (R & D Systems). Leptin concentrations in pg/ml were determined with the DRP300 Quantikine ELISA kit (R & D Systems) and adiponectin in ng/ml with the human total adiponectin/Acrp30 Quantitkine ELISA kit (R & D Systems). Determination of interleukin-1 β, interleukin-6 and Tumor Necrosis Factor α (TNFα) levels were trialed with Quantikine ELISA kits but did not lead to reliable measurements consistently above the level of detection for each assay. All ELISA analyses were carried out in the laboratory of the School of Pharmacy and Pharmaceutical Sciences at Cardiff University.

#### APOE genotyping

Participants provided a saliva sample using the self-collection kit “Oragene-DNA (OG-500) (Genotek) for DNA extraction and APOE genotyping. APOE genotypes ɛ2, ɛ3 and ɛ4 were determined by TaqMan genotyping of single nucleotide polymorphism (SNP) rs7412 and KASP genotyping of SNP rs429358. Genotyping was successful in a total of 207 participants including 165 out of the 166 individuals that had undergone an MRI scan. The genotypic distribution of those successfully genotyped can be found in Table 1 and is comparable to the expected frequencies in the normal population (Lahiri et al., 2004). In addition, participants provided information about their family history (FH) of dementia, i.e. whether a first-grade relative (parent or sibling) was affected by LOAD, vascular dementia or Lewy body disease with dementia.

### MRI data acquisition

MRI data were acquired on a 3T MAGNETOM Prisma clinical scanner (Siemens Healthcare, Erlangen, Germany) equipped with a 32-channels receive-only head coil at CUBRIC.

#### Anatomical MRI

T_1_-weighted anatomical images were acquired with a three-dimension (3D) magnetization-prepared rapid gradient-echo (MP-RAGE) sequence with the following parameters: 256 × 256 acquisition matrix, TR = 2300 ms, TE = 3.06 ms, TI = 850ms, flip angle θ = 9°, 176 slices, 1mm slice thickness, FOV = 256 mm and acquisition time of ~ 6 min.

#### High Angular Resolution Diffusion Imaging (HARDI)

Diffusion data (2 × 2 × 2 mm voxel) were collected with a spin-echo echo-planar dual shell HARDI (Tuch et al., 2002) sequence with diffusion encoded along 90 isotropically distributed orientations (Jones et al., 1999) (30 directions at b-value = 1200 s/mm^2^ and 60 directions at b-value = 2400 s/mm^2^) and six non-diffusion weighted scans with dynamic field correction and the following parameters: TR = 9400ms, TE = 67ms, 80 slices, 2 mm slice thickness, FOV = 256 × 256 × 160 mm, GRAPPA acceleration factor = 2 and acquisition time of ~15 min.

#### Quantitative magnetization transfer weighted imaging (qMT)

An optimized 3D MT-weighted gradient-recalled-echo sequence (Cercignani and Alexander, 2006) was used to obtain magnetization transfer-weighted data with the following parameters: TR = 32 ms, TE = 2.46 ms; Gaussian MT pulses, duration t = 12.8 ms; FA = 5°; FOV = 24 cm, 2.5 × 2.5 × 2.5 mm^3^ resolution. The following off-resonance irradiation frequencies (Θ) and their corresponding saturation pulse amplitude (ΔSAT) for the 11 MT-weighted images were optimized using Cramer-Rao lower bound optimization: Θ = [1000 Hz, 1000 Hz, 2750 Hz, 2768 Hz, 2790 Hz, 2890 Hz, 1000 Hz, 1000 Hz, 12060 Hz, 47180 Hz, 56360 Hz] and their corresponding ΔSAT = [332°, 333°, 628°, 628°, 628°, 628°, 628°, 628°, 628°, 628°, 332°]. The longitudinal relaxation time, T_1_, of the system was estimated by acquiring a 3D gradient recalled echo sequence (GRE) volume with three different flip angles (θ = 3,7,15). Data for computing the static magnetic field (B_0_) were collected using two 3D GRE volumes with different echo-times (TE = 4.92 ms and 7.38 ms respectively; TR= 330ms; FOV= 240 mm; slice thickness 2.5 mm) (Jezzard and Balaban, 1995).

#### Abdominal scans

Paired single-shot in-phase (TE = 2.34 ms) and out-phase (TE = 3.4 msec) gradient echo images of the abdomen were acquired at the level of the lumbar spine segment 4 (TR = 1910 ms, TI = 1200ms, flip angle θ = 20°, 10mm slice thickness). Participants were instructed to hold their breath during the brief image acquisition to minimise movement artefacts.

## MRI data processing

The two-shell diffusion-weighted HARDI data were split and b = 1200 and 2400 s/mm^2^ data were corrected separately for distortions induced by the diffusion-weighted gradients and artifacts due to head motion with appropriate reorientation of the encoding vectors (Leemans and Jones, 2009) in ExploreDTI (Version 4.8.3) (Leemans et al., 2009). EPI-induced geometrical distortions were corrected by warping the diffusion-weighted image volumes to the T_1_–weighted anatomical images which were down-sampled to a resolution of 1.5 × 1.5 × 1.5 mm (Irfanoglu et al., 2012). After preprocessing, the Neurite Orientation Dispersion and Density (NODDI) model (Zhang et al., 2012) was fitted to the dual-shell HARDI data using fast, linear model fitting algorithms of the Accelerated Microstructure Imaging via Convex Optimization (AMICO) framework (Daducci et al., 2015) to obtain isotropic signal fraction (ISOSF), intracellular signal fraction (ICSF) and orientation dispersion index (ODI) maps (Figure 2).

MT-weighted GRE volumes for each participant were co-registered to the MT-volume with the most contrast using a rigid body (6 degrees of freedom) registration to correct for inter-scan motion using Elastix (Klein et al., 2010). The 11 MT-weighted GRE images and T1-maps were modelled by the two pool Ramani’s pulsed MT approximation (Ramani et al., 2002). This approximation provided maps of the macromolecular proton fraction (MPF) and the forward exchange rate *k*_*f*_ MPF maps were thresholded to an upper intensity limit of 0.3 and *k*_*f*_ maps to an upper limit of 3 using the FMRIB’s fslmaths imaging calculator to remove voxels with noise-only data.

All image modality maps and region of interest masks were spatially aligned to the T_1_-weighted anatomical volume as reference image with linear affine registration (12 degrees of freedom) using FMRIB’s Linear Image Registration Tool (FLIRT).

## Tractography

The RESDORE algorithm (Parker et al., 2013) was applied to identify outliers, followed by whole brain tractography with the damped Richardson-Lucy algorithm (dRL) (Dell’acqua et al., 2010) on the 60 direction, b = 2400 s/mm^2^ HARDI data for each dataset in single-subject native space using in house software (Parker, 2014) coded in MATLAB (the MathWorks, Natick, MA). To reconstruct fibre tracts, dRL fibre orientation density functions (fODFs) were estimated at the centre of each image voxel. Seed points were positioned at the vertices of a 2×2×2 mm grid superimposed over the image. The tracking algorithm interpolated local fODF estimates at each seed point and then propagated 0.5mm along orientations of each fODF lobe above a threshold on peak amplitude of 0.05. Individual streamlines were subsequently propagated by interpolating the fODF at their new location and propagating 0.5mm along the minimally subtending fODF peak. This process was repeated until the minimally subtending peak magnitude fell below 0.05 or the change of direction between successive 0.5mm steps exceeded an angle of 45°. Tracking was then repeated in the opposite direction from the initial seed point. Streamlines whose lengths were outside a range of 10mm to 500mm were discarded.

The fornix, PHC, CST and UF pathways were reconstructed with an in-house automated segmentation method based on principal component analysis (PCA) of streamline shape (Parker et al., 2015). This procedure involves the manual reconstruction of a set of tracts that are then used to train a PCA model of candidate streamline shape and location. Twenty datasets were randomly selected as training data. Tracts were reconstructed by manually applying waypoint region of interest (ROI) gates (“AND”, “OR” and “NOT” gates following Boolean logic) to isolate specific tracts from the whole brain tractography data. ROIs were placed in HARDI data native space on colour-coded fiber orientation maps (Pajevic & Pierpaoli, 1999) in ExploreDTI following previously published protocols (Metzler-Baddeley et al., 2013; Metzler-Baddeley et al., 2012a; Metzler-Baddeley et al., 2011; Metzler-Baddeley et al., 2012c). The trained PCA shape models were then applied to all datasets: candidate streamlines were selected from the whole volume tractography as those bridging the gap between estimated end points of the candidate tracts. Spurious streamlines were excluded by means of a shape comparison with the trained PCA model. All automatic tract reconstructions underwent quality control through visual inspection and any remaining spurious fibers that were not consistent with the tract anatomy were removed from the reconstruction where necessary.

## Whole brain white matter and hippocampal segmentation

Whole brain white matter, left and right whole hippocampus masks were automatically segmented from T_1_- weighted images with the Freesurfer image analysis suite (version 5.3), which is documented online (https://surfer.nmr.mgh.harvard.edu/). Whole brain white matter masks were thresh-holded to exclude ventricle CSF spaces from the mask.

### Abdominal subcutaneous and visceral fat segmentation

All images were visually inspected for motion artefacts and in- and out-phase alignment in MRIcron (Rorden et al., 2007). Pure fat signal images were created with the fslmaths tool from the FSL analysis library (Jenkinson et al., 2012; Smith et al., 2004) by subtracting out-phase images of the water signal from in-phase images that contained signals from both fat and water (Dixon, 1984). Subcutaneous and visceral fat regions were then manually segmented from fat-only images in fslview.

Subcutaneous fat was defined as fat tissue exterior to the abdominal wall (see Figure 1c). Visceral fat regions were isolated by removing subcutaneous fat, muscle tissue (left and right psoas muscles; left and right internal and external oblique muscles; left and right transversus abdominis muscles; left and right rectus abdominus muscles), and the spinal disc from the images (Figure 1c). Subcutaneous and visceral masks were thresholded with an intensity of 2 to ensure that only fat tissue was included in the masks. Finally, subcutaneous and visceral fat volume fractions were obtained by dividing subcutaneous and visceral fat volumes by the total abdominal fat volume.

### Statistical analyses

Statistical analyses were conducted in SPSS version 24 (IBM, 2011) and the PROCESS computational tool for mediation analysis (Hayes, 2012). All data were inspected for assumptions of normal distribution and variance heterogeneity. Plasma adipokines, CRP and IL-6 were log-transformed to correct for skew. Multiple-comparisons-related Type 1 errors were corrected with a 5% False Discovery Rate (FDR) using Benjamini-Hochberg adjusted p-values. All p-values were two-tailed. Partial Eta^2^ (ηp^2^) and correlation coefficients are reported as indices of effect sizes. Exploratory Principal Component Analysis (PCA) was used for the purpose of assessing data dimensionality and of reducing data complexity. A PCA procedure with orthogonal Varimax rotation of the component matrix that used the Kaiser criterion of including all components with an eigenvalue of >1(IBM, 2011) was employed. Cattell’s scree plot (Cattell, 1952) and component loadings were inspected with regard to their interpretability. Loadings that exceed a value of 0.5 were considered as significant.

Omnibus multivariate regression analysis was conducted to test for the relationships between the four body composition/adiposity metrics and demographic variables (age, sex, years of education), health-related variables (alcohol consumption, blood pressure, physical activity, plasma adipokines) and genetic risk of LOAD (*APOE* genotype, FH). Post-hoc group comparisons were conducted with independent t-tests. Pearson correlations coefficients were calculated between body composition/adiposity metrics and brain measurements. Post-hoc t-test and Pearson correlations were FDR corrected. These were followed up by partial Spearman rho correlations coefficients to explore the contribution of any confounding variables identified in the multivariate omnibus regression analysis and by mediation analyses to explore for the contribution of systemic inflammation markers to the observed brain-obesity relationships.

### Missing data

Four participants did not complete the 90 minutes MRI scanning session due to claustrophobia, and qMT and abdominal MRI data are missing for these individuals. The abdominal scans were acquired at the end of the MR session and thirty-two abdominal datasets had to be excluded from the analyses due motion artefacts and due to participants holding their breath at different points of the breathing cycle during in and out-phase image acquisition, i.e. at the end of an exhalation in one image and at the end of an inhalation in the other. This meant that the image pairs were not spatially aligned and hence subcutaneous and visceral fat regions could not be reliably delineated. Furthermore, bloods could not be drawn or analysed for 18 participants. For one participant *APOE* could not be genotyped from the saliva sample and two participants did not know their family history of dementia.

## Results

### Cross-correlations between obesity metrics

Individual differences in BMI correlated positively with differences in the subcutVF [r(130) = 0.4, p < 0.001] and with differences in the WHR [r(166) = 0.21, P = 0.006] but not with differences in viscVF (p = 0.29). Differences in WHR correlated positively with differences in viscVF [r(130) = 0.28, p = 0.001] but not with subcVF (p = 0.7). There was a negative correlation between differences in viscVF and subcutVF [r(130) = −0.42, p < 0.001].

### Multivariate regression analysis exploring the relationship between body composition/adiposity metrics and demographic, health and genetic variables

Effects of demographic variables (age, sex, years of education), health-related variables (alcohol consumption, systolic and diastolic blood pressure, physical activity, plasma leptin, adiponectin, CRP, interleukin-8) and genetic risk of LOAD (*APOE* genotype, FH) on the four body composition/adiposity metrics (BMI, WHR, subcVF and viscVF) were tested with multivariate regression analysis. Omnibus effects were followed up by FDR corrected *post-hoc* comparisons.

The omnibus analysis revealed significant effects of age [F(4, 84) = 3.4, p = 0.013; ηp^2^ = 0.14], sex [F(4, 84) = 9.1, p < 0.001, ηp^2^ = 0.3], adiponectin [F(4, 84) = 2.5, p = 0.046, ηp^2^ = 0.11] and leptin [F(4, 84) = 12.2, p < 0.001, ηp^2^ = 0.37] on the four body composition/adiposity metrics.

**Figure 2.**
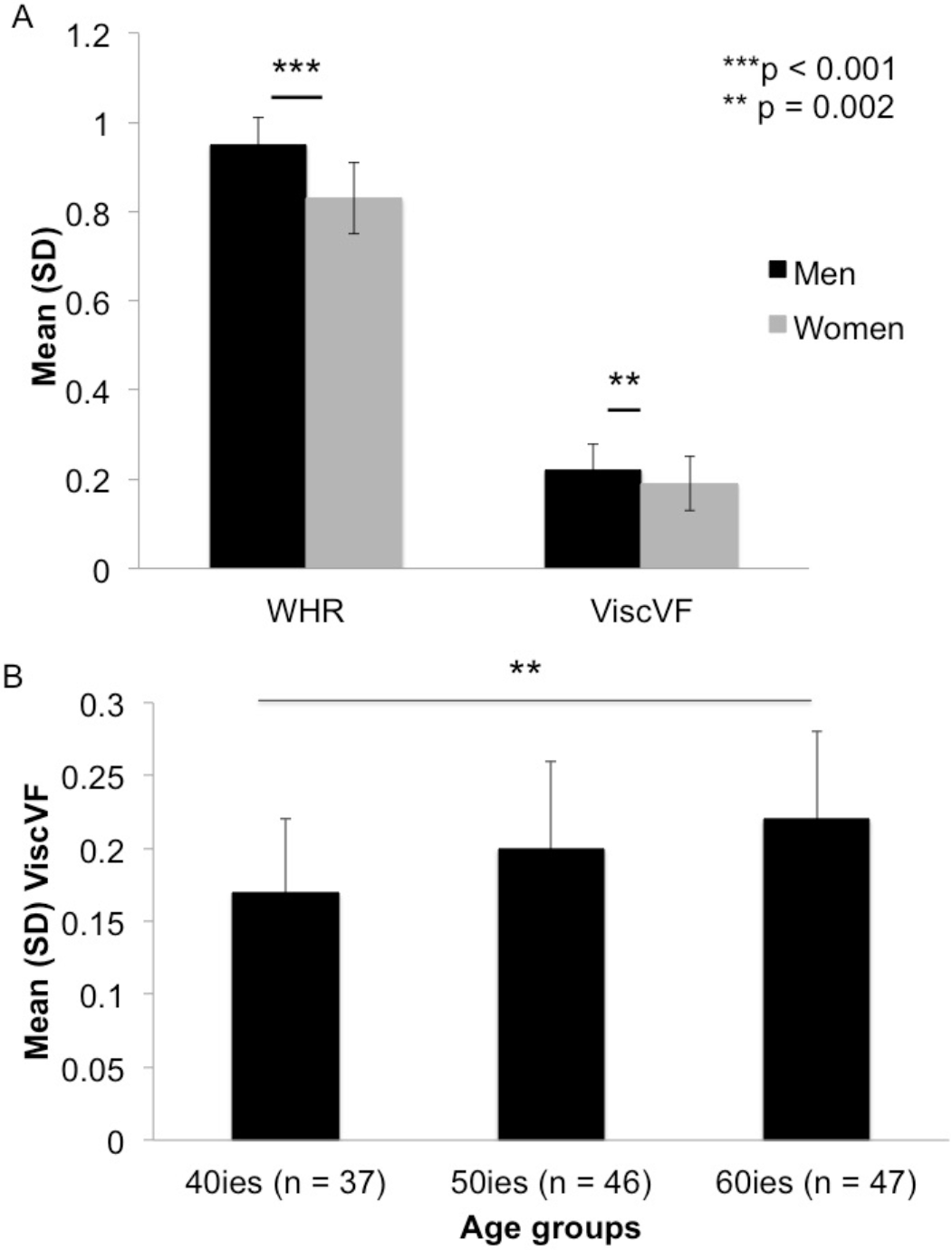
A) Men (black) showed larger Waist-to-Hip Ratios (WHR) and Visceral fat Volume Fractions (viscVF) than women (grey). B) Older individuals over 60 years of age had larger viscVF than younger individuals in their 40ies. ** FDR-corrected p-values.

**Figure 3.**
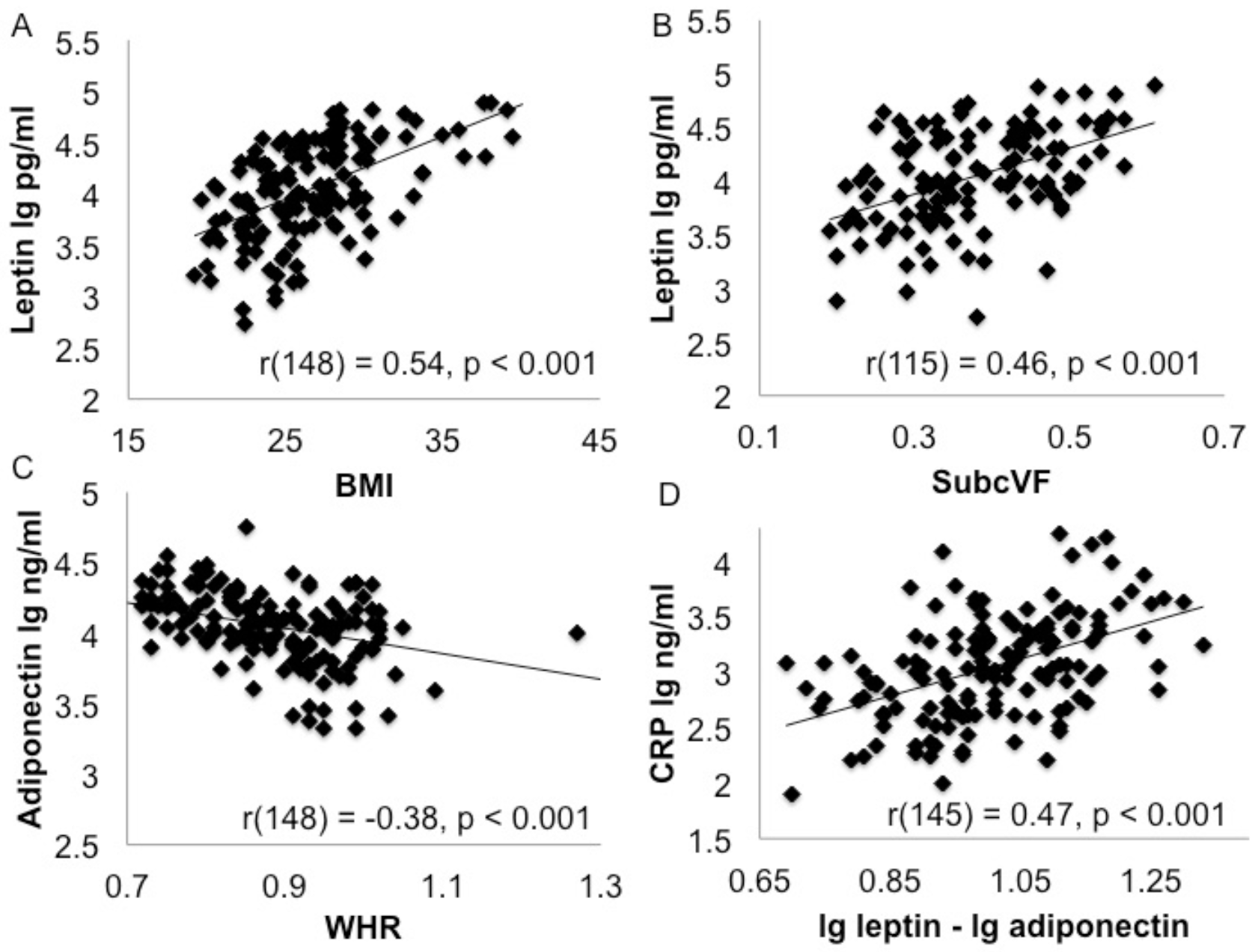
A) Individual differences in plasma leptin concentrations were positively correlated with differences in the body mass index (BMI) and b) differences in subcutaneous fat volume fraction (subcVF). C) Differences in plasma adiponectin concentrations correlated negatively with the waist hip ratio (WHR) and D) differences in the leptin/adiponectin ratio were positively correlated with differences in plasma concentrations of C-Reactive Protein (CRP).

### Post-hoc

comparisons showed that men had higher WHR [t(164) = 5.5, p < 0.001] and viscVF [t(128) = 3.1, p = 0.002] than women (Figure 2a). Older individuals over 60 had larger viscVF than younger individuals in their 40ies [t(82) = 3.24, p = 0.002] (Figure 2b). Leptin was positively correlated with BMI [r(148) = 0.54, p < 0.001] (Figure 3a) and subcutVF [r(115) = 0.46, p < 0.001] (Figure 3b), and adiponectin was negatively correlated with WHR [r(148) = −0.38, p < 0.001] (Figure 3c).

### Dimensionality of microstructural indices in white matter and hippocampal gray matter

PCA explored the dimensionality of the microstructural indices in all regions of interest. This was done separately for white and gray matter as the two tissue types differ in their microstructural organisation, i.e., white matter consists of relatively aligned axons and glia, whilst gray matter comprises a more complex microstructure of cell bodies, dendrites, synapses and glia. PCA was also employed to reduce the complexity of the microstructural data [40 measurements for white matter and 10 for the hippocampus) for subsequent correlation analyses with the body composition/adiposity metrics.

Table 2 summarizes the component loadings for the extracted WM components. PCA resulted in five components that all exceeded an eigenvalue of 2 and explained together 65.4% of the data variation. The first component had high loadings of ISOSF and ODI, the second of *k*_*f*_, the third of MPF, and the fourth of ICSF in all regions. The fifth component had loadings from ISOSF, MPF and *k*_*f*_, of the fornix only. PCA of microstructural metrics from left and right hippocampi extracted four components that accounted for 75% of the data. These components were: 1^st^ ODI and MPF, 2^nd^ ISOSF, 3^rd^ ICSF, and 4^th^ *k*_*f*_ from left and right hippocampi (Table 3).

**Table 2.** Rotated component matrix of the principal component analysis of the white matter microstructural indices ^a^

| ROI | WM index | Components |  |  |  |  |
| --- | --- | --- | --- | --- | --- | --- |
| | | ISOSF/ODI | $k_f$ | MPF | ICSF | Fornix |
| Fornix | ICSF | -0.201 | -0.039 | -0.081 | 0.497 | 0.431 |
|  | ISOSF | -0.045 | 0.012 | -0.015 | 0.039 | **-0.93** |
|  | ODI | **0.622** | -0.041 | -0.027 | -0.122 | -0.364 |
|  | MPF | -0.114 | 0.041 | 0.458 | 0.049 | **0.782** |
| | $k_f$ | -0.09 | **0.525** | -0.014 | 0.008 | **0.757** |
| Left PHC | ICSF | 0.264 | 0.107 | 0.101 | **0.792** | -0.191 |
|  | ISOSF | **0.781** | -0.045 | 0.01 | 0.11 | -0.042 |
|  | ODI | **0.641** | -0.251 | -0.091 | -0.287 | 0.123 |
|  | MPF | 0.124 | 0.276 | **0.552** | 0.347 | 0.005 |
| | $k_f$ | 0.09 | **0.814** | -0.048 | 0.18 | -0.004 |
| Right PHC | ICSF | 0.116 | 0.054 | 0.225 | **0.77** | -0.233 |
|  | ISOSF | **0.752** | -0.008 | 0.09 | -0.08 | -0.081 |
|  | ODI | **0.621** | -0.086 | -0.155 | -0.285 | 0.206 |
|  | MPF | 0.049 | 0.149 | **0.701** | 0.265 | -0.01 |
| | $k_f$ | 0.007 | **0.82** | 0.026 | 0.156 | 0.107 |
| Left UF | ICSF | -0.043 | 0.075 | 0.377 | **0.743** | 0.007 |
|  | ISOSF | **0.728** | -0.062 | 0.234 | -0.083 | -0.029 |
|  | ODI | **0.831** | -0.054 | -0.066 | -0.103 | 0.028 |
|  | MPF | 0.018 | 0.262 | **0.76** | 0.182 | 0.04 |
| | $k_f$ | -0.154 | **0.814** | 0.178 | 0.108 | -0.054 |
| Right UF | ICSF | -0.039 | 0.09 | 0.427 | **0.647** | 0.089 |
|  | ISOSF | **0.642** | -0.021 | 0.226 | -0.28 | 0.066 |
|  | ODI | **0.781** | -0.03 | -0.025 | -0.154 | 0.039 |
|  | MPF | -0.027 | 0.204 | **0.772** | 0.086 | 0.002 |
| | $k_f$ | 0.04 | **0.752** | 0.351 | 0.005 | -0.043 |
| Left CST | ICSF | -0.222 | 0.02 | 0.031 | **0.66** | 0.085 |
|  | ISOSF | **0.742** | -0.087 | -0.032 | 0.077 | -0.088 |
|  | ODI | **0.797** | 0.029 | 0.022 | 0.036 | 0.097 |
|  | MPF | -0.067 | -0.045 | **0.83** | 0.132 | 0.042 |
| | $k_f$ | -0.155 | **0.818** | 0.107 | -0.088 | 0.073 |
| Right CST | ICSF | -0.271 | 0.049 | 0.147 | **0.56** | 0.079 |
|  | ISOSF | **0.716** | -0.116 | -0.127 | 0.115 | -0.132 |
|  | *ODI* | **0.826** | 0.026 | -0.053 | 0.064 | 0.065 |
|  | *MPF* | -0.114 | -0.024 | **0.85** | 0.077 | -0.015 |
|  | *k<sub>f</sub>* | -0.167 | **0.802** | 0.169 | -0.067 | 0.069 |
| *Total WM* | *ICSF* | -0.298 | 0.001 | 0.267 | **0.701** | -0.333 |
|  | *ISOSF* | **0.752** | -0.037 | -0.105 | -0.147 | 0.201 |
|  | *ODI* | **0.527** | 0.06 | -0.186 | 0.005 | -0.195 |
|  | *MPF* | -0.065 | 0.1 | **0.812** | 0.196 | 0.235 |
|  | *k<sub>f</sub>* | -0.095 | **0.928** | 0.051 | 0.047 | 0.062 |

Loadings > 0.5 are highlighted in bold. Abbreviations: CST= corticospinal
tract, ICSF = intracellular signal fraction, ISOSF = isotropic signal fraction, *k<sub>f</sub>*
= forward exchange rate, MPF = macromolecular proton fraction, ODI =
orientation dispersion index, PHC = parahippocampal cingulum, UF =
uncinate fasciculus, WM = white matter.
<sup>a</sup> Rotation method: Varimax with Kaiser normalization.

**Table 3.** Rotated component matrix of the principal component analysis of the hippocampal microstructural indices^a^

| *ROI* | *GM index* | **Components** |  |  |  |
| --- | --- | --- | --- | --- | --- |
|  |  | **ODI/MPF** | **ISOSF** | **ICSF** | ***k<sub>f</sub>*** |
| *Left HC* | *ICSF* | 0.047 | -0.058 | **0.941** | 0.066 |
|  | *ISOSF* | 0.104 | **0.877** | -0.015 | -0.174 |
|  | *ODI* | **0.834** | 0.106 | 0.005 | -0.246 |
|  | *MPF* | **-0.537** | -0.378 | 0.236 | -0.195 |
|  | *k<sub>f</sub>* | -0.044 | -0.118 | -0.02 | **0.859** |
| *Right HC* | *ICSF* | -0.191 | -0.078 | **0.901** | -0.033 |
|  | *ISOSF* | 0.153 | **0.846** | -0.077 | -0.262 |
|  | *ODI* | **0.863** | 0.025 | 0.005 | -0.217 |
|  | *MPF* | **-0.526** | -0.437 | 0.218 | -0.115 |
|  | *k<sub>f</sub>* | -0.194 | -0.189 | 0.05 | **0.807** |

Loadings > 0.5 are highlighted in bold. Abbreviations: GM = gray matter, HC = hippocampus, ICSF = intracellular signal fraction, ISOSF = isotropic signal fraction, *k<sub>f</sub>* = forward exchange rate, MPF = macromolecular proton fraction, ODI = orientation dispersion index.
<sup>a</sup> Rotation method: Varimax with Kaiser normalization.

### Correlations between body composition/adiposity metrics and white and gray matter microstructure

Pearson correlation coefficients were calculated between BMI, WHR, subcVF, and viscVF and the following brain components: WM ODI/ISOSF, WM ICSF, WM MPF, WM *k*_*f*_ hippocampal ODI/MPF, hippocampal ISOSF, hippocampal ICSF and hippocampal *k*_*f*_. As fornix was a separate component in the PCA with high loadings from fornix MPF, *k*_*f*_, and ISOSF, these variables were also included in the analyses. WHR was positively associated with fornix ISOSF [r(166) = 0.29, p < 0.001] and the hippocampal ISOSF component [r(158) = 0.29, p < 0.001], and negatively with fornix MPF [r(162) = −0.3, p < 0.001] and fornix *k*_*f*_ [r(162) = −0.28, p < 0.001]. ViscVF correlated negatively with fornix MPF [r(129) = −0.3, p = 0.001 and fornix *k*_*f*_ [r(129) = −0.33, p < 0.001] and positively with fornix ISOSF [r(130) = 0.25, p = 0.004].

**Figure 4.**
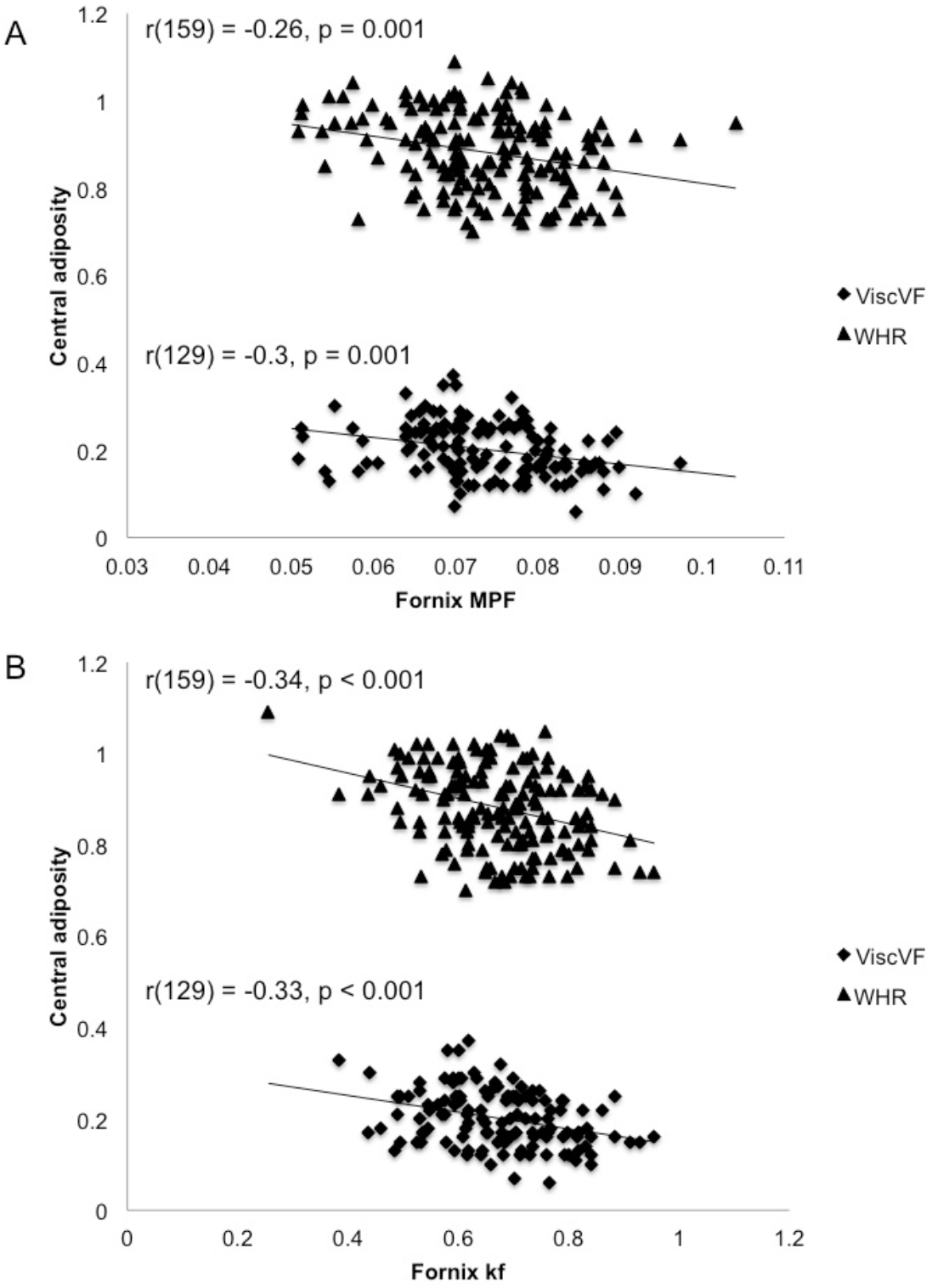
A) Fornix macromolecular proton fraction (MPF) was negatively correlated with Waist-to-Hip Ration (WHR) and visceral volume fraction (viscVF). B) Fornix forward exchange rate *k*_*f*_ was negatively associated with WHR and viscVF. All p-values were FDR corrected. Note that three extreme outliers in the WHR variable, that deviated more than three standard deviations from the regression slope, were removed from the scatterplot for display purposes, but their removal did not alter the results.

Controlling for the variables that showed significant omnibus effects in the multivariate regression analysis above, i.e., age, sex, leptin and adiponectin with partial Spearman rho correlation, removed all correlations with fornix ISOSF (p = 0.15 and 0.35) and hippocampal ISOSF (p=0.18). However, the correlations between WHR and fornix *k*_*f*_ [r(108) = −0.26, p = 0.007], between WHR and fornix MPF [r(108) = −0.21, p = 0.03], and between viscVF and fornix MPF [r(108) = −0.22, p = 0.02] remained significant (Figure 4).

### Mediation analysis testing for the contribution of systemic inflammation

Mediation analyses testing for the contributing effects of individual differences in CRP and interleukin-8 plasma concentrations as well as in the leptin/adiponectin ratio (difference between log10 leptin and log10 adiponectin plasma concentrations) (Lopez-Jaramillo et al., 2014) to the WHR/viscVF- fornix MPF/*k*_*f*_ relationships were carried out. Differences in the leptin/adiponectin ratio contributed significantly to the correlations between fornix MPF and WHR (t = 3.1, p = 0.002), and fornix MPF and viscVLF (t= 3.3, p = 0.001) but not to the correlations with fornix *k*_*f*_. CRP and interleukin-8 did not significantly contribute, but CRP was correlated positively with the leptin/adiponectin ratio [r(145) = 0.47, p < 0.001].

## Discussion

Midlife obesity is a risk factor of LOAD, but the biological mechanisms underpinning this link remain poorly understood. Both conditions are associated with systemic inflammation, and it is increasingly recognised that microglia-mediated immune responses play an important role in LOAD (Dansokho and Heneka, 2018; Heneka et al., 2015; Sarlus and Heneka, 2017; Tejera and Heneka, 2016). It has therefore been proposed that obesity induced gut dysbiosis may trigger micro-glia mediated neuroinflammation that in turn may contribute to the development of LOAD pathology (Bartzokis, 2011; Sochocka et al., 2018; Venegas et al., 2017). If that was the case, one might expect adverse effects of obesity-related neuroinflammation to manifest in brain regions involved in LOAD. Furthermore, as the pathological processes leading to LOAD are likely to accumulate over many years (Jack et al., 2013), it may be possible to identify such brain tissue changes in cognitively healthy individuals prior to the onset of any memory symptoms.

Here we tested this hypothesis by studying the impact of central obesity on MRI indices of white matter microstructure in limbic regions in a cohort of cognitively healthy individuals (38 – 71 years of age) that were well characterised with regards to their lifestyle and genetic risk of LOAD (Table 1). Consistent with the above hypothesis, we observed central obesity related reductions in myelin-sensitive metrics of fornix white matter that were not explained by differences in demographic, health or genetic risk related variables.

More specifically, individual differences in body adiposity were measured with BMI, WHR, and with MRI estimates of abdominal fat distribution i.e. with visceral and subcutaneous fat volume fractions. In addition, tissue microstructure was assessed with quantitative MRI indices that allowed the separate investigation of white matter properties related to myelin/ tissue metabolism and axon density.

Correlation analyses revealed positive correlations between WHR and viscVF but not with subcutVF. In contrast, BMI correlated positively with subcutVF but not with viscVF. Whilst BMI and WHR were positively correlated, viscVF and subcVF correlated negatively with each other. From this pattern of cross-correlations, it is clear that BMI and WHR captured quite different fat distributions in our sample. Indeed, negative correlations with white matter microstructure were only observed for WHR and viscVF, but not for BMI or subcVF. These results are consistent with previous findings of visceral but not subcutaneous fat being associated with an increased risk of metabolic syndrome and mortality (Koster et al., 2015; Koster and Schaap, 2015; Koster et al., 2010) as well as with reduced brain volume (Debette et al., 2010).

The observed pattern of correlations also implies that the direct comparison of results across studies with different measures of body obesity/distribution may be difficult. This observation may explain discrepant findings in the literature, as some studies reported beneficial effects of larger BMI or waist circumference on white matter microstructure (Birdsill et al., 2017), whilst other studies reported adverse effects Kullmann et al., 2015; Ronan et al., 2016). Previously we reported significant correlations between BMI and fornix white matter microstructure in a smaller group of older adults (Metzler-Baddeley et al., 2013), a result that was not replicated here. At first glance these findings seem at odds with the present study. However, the first study investigated white matter microstructure with DTI indices of fractional anisotropy (FA), mean, axial, and radial diffusivity, and observed positive correlations between BMI and the diffusivities, but no correlation with FA. DTI indices are nonspecific metrics of white matter microstructure that are affected by changes in biological white matter properties as well as by their geometrical and organisational architecture (Beaulieu and Allen, 1994; De Santis et al., 2014). Hence it is not possible to interpret BMI-related increases in fornix diffusivities in terms of differences in myelin or axon density. Furthermore, participants in the first study were older adults with BMI levels from within the normal to overweight range (53-93 years of age, Mean_age_ = 68, Mean_BMI_ =24.9, n = 38), whilst the majority of middle-aged participants in this study were overweight (Mean_BMI_ =27), with 20% falling within the obese category. As we proposed in the previous paper, the observed relationship between BMI and fornix microstructure may not relate to mechanisms underpinning obesity, but may rather reflect some functional properties of the fornix within hippocampal-hypothalamic-prefrontal food control networks (Metzler-Baddeley et al., 2013).

The results of the PCA for the white matter microstructural indices were consistent with our assumption that MPF, *k_f_* and ICSF provide estimates of different white matter tissue properties, i.e. of apparent myelin, inflammation-related tissue metabolism, and apparent axon density. ODI and ISOSF, however, were jointly loading on one component, suggesting that in our dataset they captured overlapping microstructural features. Exploratory PCA of the microstructural indices in hippocampal gray matter revealed separate components for *k*_*f*_, ICSF, and ISOSF but here ODI and MPF were found to load jointly on one component. This suggests that qMT and NODDI indices may not be directly comparable across white and gray matter. As they have primarily been validated in white matter (Sled, 2017; Zhang et al., 2012), their interpretation in gray matter remains speculative and requires histological validation.

In white matter, however, we only observed negative correlations between WHR/viscVF and the qMT metrics of MPF and *k*_*f*_ but not with ICSF, suggesting that central obesity impairs apparent myelin glia properties rather than apparent axon loss. Furthermore, the correlations between WHR and viscVF and myelin/inflammation-sensitive metrics were specifically observed in the fornix tract but not for microstructural components across the other white matter regions of interest. As the fornix is known to be impaired in Mild Cognitive Impairment and early LOAD (Metzler-Baddeley et al., 2012b; Oishi and Lyketsos, 2014; Oishi et al., 2012; Plowey and Ziskin, 2016; Yu et al., 2017), this pattern of results is consistent with the view that visceral fat may be associated with processes that have adverse effects on limbic areas involved in LOAD. Indeed, animal studies have shown that diet-induced obesity can trigger microglia-mediated inflammation in the hippocampus that impairs synaptic functioning and spatial memory (Hao et al., 2016). In the present study, we did not observe specific effects on hippocampal microstructure. However, for the reasons outlined above, one may expect qMT metrics to be more sensitive to myelin damage in white rather than in gray matter, especially as we were studying a sample of cognitively healthy individuals.

Omnibus regression analysis identified effects of sex, age, leptin and adiponectin on the obesity measurements. Controlling for these variables reduced the size but did not fully remove the correlations between WHR and fornix *k*_*f*_, or between viscVF and fornix MPF, suggesting that these effects were not simply reflecting the observed age and sex differences in WHR and viscVF.

In contrast, education, blood pressure, weekly alcohol consumption, physical exercise, *APOE* genotype and family history of dementia had no effect on interindividual differences in obesity measures in our sample. There was also no direct relationship between obesity measures and plasma CRP and interleukin-8 concentrations. However, differences in the leptin/adiponectin ratio correlated positively with differences in CRP concentrations (Figure 3) and contributed significantly to the correlations between WHR/viscVF and fornix MPF. In addition, adiponectin, a hormone with anti-inflammatory properties (Graßmann et al., 2017) was negatively correlated with WHR (Figure 3). Together, this pattern of results suggests that an obesity-related shift in the ratio between leptin and adiponectin, due to increases in leptin and reductions in adiponectin (Figure 3), determines the net effect on inflammatory mediators such as CRP. Thus it is possible, that WHR-related reductions in adiponectin, by means of exacerbating neuroinflammation (Ouchi and Walsh, 2012), may have contributed to WHR-related myelin damage in the fornix.

The fact that we did not observe any direct correlations between CRP and interleukin-8 concentrations and WHR/viscVF or fornix microstructure also needs to be seen within the context that in our cohort of middle-aged healthy participants without clinical inflammatory conditions, plasma concentrations of inflammatory markers such as interleukin-1β, interleukin-6 and TNFα were generally below the limit of detection of the assays, and we had to employ high sensitivity ELISA kits to measure CRP and Interleukin 8. Thus, the lack of direct correlations with obesity and microstructural indices may also partly be due to the very low levels of inflammatory markers in our cohort of healthy individuals.

To summarise, the present study demonstrated that central obesity, notably abdominal visceral fat accumulation, was associated with reductions in qMT indices of apparent myelin and tissue metabolism in a sample of cognitively healthy individuals at midlife and early older age. The precise mechanisms underpinning this relationship are likely to be complex. However, our results suggest that obesity-related shifts in the leptin/adiponectin ratio, that was positively correlated with the systemic inflammation marker CRP, contribute to obesity related reductions of myelin-sensitive metrics in the fornix. Overall, these results are consistent with the view that obesity-related changes in immune responses, and associated white matter glia changes may contribute to the link between midlife obesity and LOAD.

## Acknowledgements

This research was funded by a Research Fellowship to CM-B from the Alzheimer’s Society and the BRACE Alzheimer’s Charity (RES19962). DKJ is supported by a Wellcome Trust Investigator Award (096646/Z/11/Z) and a Wellcome Trust Strategic Award (104943/Z/14/Z). We would like to thank Peter Hobden and Sonya Foley-Bozorgzad for their assistance with MRI data acquisition and processing, Rosie Dwyer, Samantha Collins, Abbie Stark, and Emma Blenkinsop for their assistance with the collection and scoring of the cognitive and health data, and Rhodri Thomas for his assistance with the *APOE* genotyping of the saliva samples. The authors declare no competing financial or non-financial interests.

**Author contributions**
CM-B is the PI of the study and is responsible for the conceptualization and data acquisition and analyses of the study. CM-B has also written the manuscript. JPM and EL were responsible for participant recruitment, data acquisition and MRI data processing. RS was responsible for the *APOE* genotyping. FF and JE have prepared the qMT and diffusion MRI protocols and have helped with MRI data processing. AK-M and FG carried out the manual segmentations of the abdominal fat volume regions. RJB was involved in the conceptualisation and has advised on statistical data analysis. EK and BE were responsible for the ELISA serum analyses. DKJ provided feedback on the study design and manuscript.

